# Confinement considerations for CRISPR toxin-antidote gene drive systems

**DOI:** 10.64898/2026.08.03.742636

**Authors:** Ziqian Xu, Yuna I. Cho, Xinyue Zhang, Jackson Champer

**Author notes:** equal contribution (Ziqian Xu, Yuna I. Cho, Xinyue Zhang).

## Abstract

Gene drives are potentially powerful tools, able to spread throughout target populations. They could be used to modify or suppress disease vectors, invasive species, and agricultural pests. Yet, in many scenarios, confinement of the drive to only a target population is required. Several types of drives are capable of this, among the most promising of which are CRISPR toxin-antidote drives. Though showing good performance in simple models, such drives have not been thoroughly assessed in spatially structured populations. Here we evaluate three modification drive variants with varying levels of confinement. We find that Toxin-Antidote Recessive Embryo (TARE) drive and 2-locus TARE drives can usually spread in connected populations or from a single sufficiently large release. However, they can still be stopped by a migration corridor or by a sufficiently high population density gradient. 1-locus 2-drive TARE, on the other hand, will not be able to spread outside of release areas and indeed will often retreat in the face of wild-type alleles. When these drives arrive at a point source such as a port, only the standard TARE drive has a significant chance of establishing if releases occur at sufficiently high frequency and quantity. However, in models with parental care among a limited number of offspring, TARE drives become more invasive, with lower introduction thresholds. Overall, we find that spatial and other ecological factors can substantially affect the outcome of a confined CRISPR toxin-antidote drive release.

## Introduction

Gene drive is a phenomenon of biased inheritance where a certain genetic element or allele is inherited at an increased rate^1–7^. Currently, vector-borne diseases, agricultural pests, and invasive species are major problems worldwide. Vector-borne diseases, such as malaria, dengue, West Nile fever, Zika, and Lyme disease, represent 17% of all infectious diseases, causing more than 700,000 deaths annually^8^. According to the USDA, pests reduce global crop production by 20% to 40% annually^9^. Invasive species are the second most common cause of species extinctions in the past 500 years^10^. In many of these cases, gene drive is a potential solution^11,12^.

There are two general types of gene drives classified by their effect: population suppression and population modification^1–7^. Modification drives can be used to add new genes or modify existing mosquito genes to make mosquitoes refractory to *Plasmodium*, the malaria parasite, or to other diseases. In contrast, suppression drives can be used to eliminate the whole population by disrupting a vital gene to generate phenotypes such as recessive female sterility or even lethality. Both types of gene drives have been demonstrated in a variety of species^1–7^.

However, well-studied homing type gene drives that copy themselves are powerful tools, potentially affecting a whole species. They can easily spread after an introduction of just a few individuals, which can be problematic in some scenarios^13^. For example, sociopolitical reasons may require that spread of a gene drive be confined to a particular region. Targeting of invasive species and agricultural pests should also be conducted only where the species are pests. Finally, hybrids may result in gene drive spread outside the target species, which would only be desirable when other closely related species are also targets. Thus, unconfined homing drives and similar systems may not be suitable for addressing situations featuring such issues.

Gene drive confinement can be an answer to this challenge. Specifically, several gene drives are based on an introduction threshold^1–3,14–17^. If their frequency in a population is below the threshold, the drive will decline toward elimination. Only if the drive frequency is above the threshold can it increase its overall frequency in the population. There are many types of confined drives, but CRISPR toxin-antidote drives, in particular, have recently garnered much attention. They target and disrupt an essential gene with CRISPR, but also provide a rescue element for the target that cannot be cleaved. Thus, wild-type alleles are removed, but the drive persists, increasing in frequency. So far, several similar variants of these drives, such as Toxin-Antidote Recessive Embryo (TARE)^18–21^ and Cleave and Rescue (ClvR)^22–25^ have been tested in flies, mosquitoes^26^, and some variants even in model plants^27,28^ (though the confinement properties for plant drives are based on species-specific factors^29^). While basic forms only have an introduction threshold with fitness costs^30,31^, higher introduction thresholds for greater confinement are possible. For this, underdominance variants are most suitable^21,32^. Overall, these gene drive strategies are potentially both flexible and limited in their spread.

However, previous studies have indicated the confinement of gene drives is more complicated in models with continuous space^33–38^. While networks of linked demes^39,40^ may be suitable in some situations, continuous space may be more representative of many real-world scenarios where individuals move over a small part of a larger landscape. In such models, allele movement usually results in some gene drive regions and some wild-type regions, with dynamic interactions between them. For example, drives with high introduction thresholds, but still below 50%, can still usually form a wave of advance in an open region, eventually spreading to the whole connected population. Yet, if released in a limited area, even drives with low thresholds may be eliminated rapidly due to influx of surrounding wild-type alleles. In some cases, geographical barriers can still successfully confine an invasive drive or allow a strongly confined drive (with an introduction threshold of over 50%) to persist^33,35^. Several variants of underdominance designs^33,36–38^ and *Wolbachia* bacteria^41–45^ (which spread with similar population dynamics) have been assessed in these models, but newer, highly promising CRISPR-based toxin-antidote drives have only been assessed in simple panmictic models^22,30,32^, for specific population suppression configurations in spatial models^31,34,35^, and for characterizing release and developing optimal release methods^46,47^.

To address this gap, in this study, we assess confinement of three representative CRISPR toxin-antidote drives in more depth. These include an invasive TARE drive, a 2-locus TARE drive representing an underdominance system, and a highly confined 1-locus 2-drive TARE system. We assess their behavior in open fields, but also several other spatial scenarios, finding parameter ranges where they invade or persist in migration corridors, density gradients, and when invading new regions from a point source. We then further consider other ecological complications that could alter the confinement level of these gene drive systems, particularly reductions in introduction thresholds in species with parental care due to greater offspring survival^48–52^. We find that successful confinement is usually possible with some variants in each considered scenario, but care must be taken to avoid overly invasive gene drive systems, necessitating the use of specific models.

## Methods

### Drive systems

We modeled three different TARE systems previously reported, which vary in configuration and introduction threshold frequencies (Figure 1).

1. Standard TARE. This is the initial TARE system introduced^20,22,30^, which comprises a single drive allele at a single genomic locus containing an essential but haplosufficient gene. The TARE drive will cut this gene and thus turn wild-type alleles into disrupted alleles at 100% efficiency in the germline. Progeny of female drive carriers will also have any wild-type alleles converted to disrupted alleles due to maternally deposited Cas9 and gRNA. Any individual that is homozygous for disrupted alleles will be nonviable. This drive has no introduction threshold when it has no fitness costs. Here, we model the TARE system with a default small fitness cost (homozygote fitness of 0.9, with multiplicative per-allele fitness), which yields an introduction threshold frequency of 4.6%.
2. 2-locus TARE. This system contains two TARE drive alleles at two (usually unlinked) loci^21,32^. Each is similar to a normal TARE system, but their gRNAs target the gene where the other drive is placed. Any individuals homozygous for disrupted alleles at either locus are nonviable. Our default model (unless otherwise specified) is a 2-locus TARE drive without fitness costs, which has an introduction threshold frequency of 18%.
3. 1-locus 2-drive TARE. This system consists of two different TARE drive alleles at the same genomic locus, each targeting a different essential but haplosufficient gene^32^. These are distant-site drives, so their target genes are each at unlinked loci. Both drives provide rescue for one of these genes, and target the gene that the other drive rescues. Because both drives are usually required for viability and they must share the same locus, more drive genotypes are nonviable. Our default model is a system with no fitness costs, which has a high introduction threshold frequency of 61%.
4. Normal allele. In some scenarios, we model a normal allele that follows Mendelian inheritance and has no special properties.

**Figure 1.**
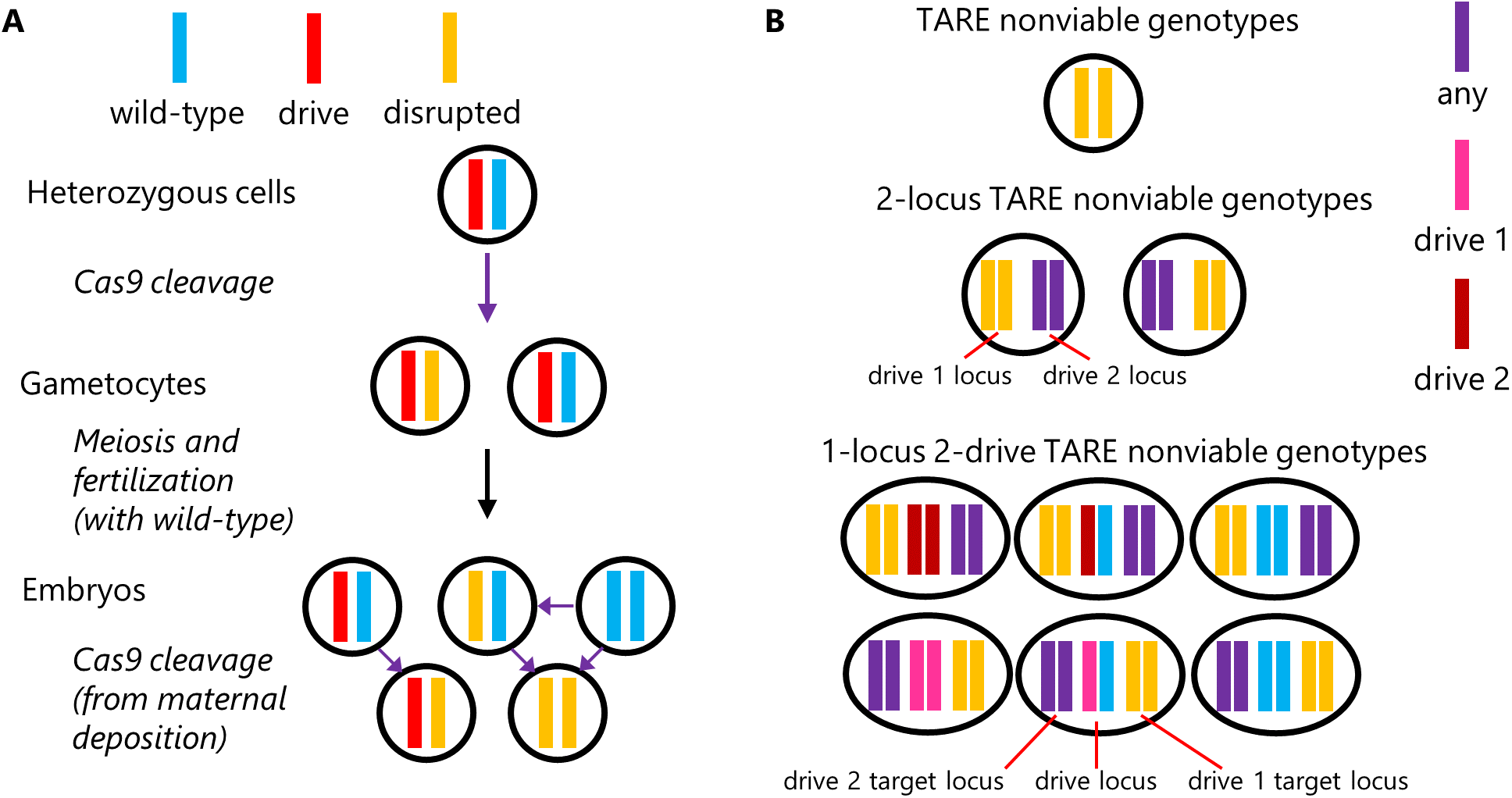
CRISPR toxin-antidote drive systems. (**A**) TARE drives target essential but haplosufficient genes, converting wild-type alleles to disrupted alleles in germline cells. Additional wild-type alleles in embryos of drive females can also be converted to disrupted alleles after cleavage by maternally deposited Cas9 and gRNA. (**B**) In standard TARE drive, the only nonviable genotype is disrupted allele homozygotes. In 2-locus TARE, individuals are nonviable if they have only disrupted alleles at either locus. Similarly, in 1-locus 2-drive TARE, individuals are nonviable if they have only disrupted alleles at either drive site and lack a drive that can provide rescue at the drive site.

### Population Model

Each of the TARE systems is implemented in an individual-based, forward-in-time population genetic simulation framework using SLiM 3^53^. Our model simulates a population of males and females with discrete, non-overlapping generations and is similar to previous studies^33,34,54^.

To obtain individuals for the next generation, each female selects a male. Drive-carrying males have a probability of being selected proportional to their fitness. In spatial models, mating is possible only within the average dispersal distance.

The number of offspring generated for each female is drawn from a binomial distribution with 50 draws (representing the maximum possible number of offspring) and a probability for each draw equal to female fitness/25. This ensures that wild-type females with a fitness of 1 and at normal population density have two offspring on average. Fitness is determined by genotype and further multiplied by a density-dependent competition factor. This factor is usually equal to 10N/(1 + 9N/K), where N is the current total population size and K is the carrying capacity (set to 50,000 by default). This Beverton-Holt model produces logistic growth. For spatial models, K is the carrying density (by default, set to produce a density of 50,000 individuals in a 1x1 two-dimensional arena or a one-dimensional arena of length 1), and N is the number of individuals within the competition radius, which is set as 0.01.

Next, offspring are generated by randomly receiving an allele from each parent. Any wild-type allele is converted to a disrupted allele if the parent also has an appropriate drive allele. Any wild-type allele in new offspring then is also converted to a disrupted allele if the mother had at least one drive allele due to maternal deposition of Cas9 and gRNA. All offspring with nonviable genotypes are removed from the population. In spatial models, each offspring is displaced from their mother in a random direction and a distance drawn from an exponential distribution with a mean equal to the dispersal value. Offspring positions are redrawn if they fall outside the arena, which consists of a 1x1 square unless otherwise specified.

### Density gradient simulations

We used one-dimensional spatial simulations with a landscape of length 1. Population density was implemented through density-dependent fecundity. For an individual at position x, adult competition was calculated from the adult interaction kernel and scaled as

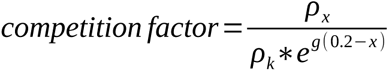

where *g* represents the density gradient factor. The local density of each position *ρ_x_* is defined as the density of individuals within a radius equal to the interaction distance. This was chosen to produce a constant population density gradient at all positions.

The expected average carrying density of individuals at normal equilibrium is

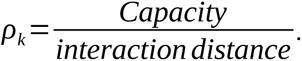

Offspring production was then multiplied by

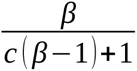

where *β* represents the low-density growth rate of the population. Thus, equilibrium density is at the even-field simulation level at position 0.2 in the density-gradient simulations and changes exponentially with position according to the density-gradient parameter. Negative density-gradient values produce increasing equilibrium density toward the right side of the habitat.

For TARE and 2-locus TARE, individuals with starting position x < 0.2 were converted to drive allele homozygotes (double homozygotes for 2-locus TARE). For 1-locus 2-drive TARE, individuals on the right side (x > 0.2) were converted to heterozygotes with both types of drive alleles. Wave speed was measured from the interpolated generation in which the fraction of individuals carrying any drive reached 0.5 in two spatial slices of width 0.1 beginning at 0.3 and 0.7, divided by the distance between those slices and normalized by dispersal. For 1-locus 2-drive TARE, this wave-speed measurement corresponds to the wild-type wave moving through the drive region.

### Density step simulations

In density-step simulations, the one-dimensional habitat was divided at x = 0.5. The right side had carrying density multiplied by DENSITY_STEP (representing the relative increase in density in the right region) relative to the left side, implemented as

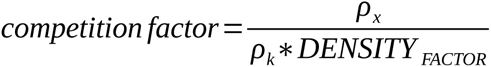

where DENSITY_FACTOR was 1 (representing the relative density of the left region) for x < 0.5 and DENSITY_STEP for x ≥ 0.5. The same Beverton-Holt density-dependent fecundity function was used to translate this competition factor into offspring number. TARE and 2-locus TARE were initialized with drive carriers on the left and wild type on the right. 1-locus 2-drive TARE was initialized in the opposite orientation, with wild type on the left and drive carriers on the higher-density right side. Outcomes were recorded as drive invasion success, drive loss/wild-type invasion success, or persistence to generation 1,000 without either outcome.

### Reproductive compensation

In some species, offspring from the same parent may directly or indirectly compete. Thus, offspring with nonviable siblings (specifically disrupted allele homozygotes formed by TARE drives in this study) may survive or be produced at higher rates. We term this “reproductive compensation.” To investigate the effect of reproductive compensation, we developed a discrete-generation, deterministic model using Excel. This model tracks genotype frequencies in each generation, assuming random mating. Fitness costs were parameterized for drive homozygotes (affecting female fertility and male mating success) with multiplicative per-allele drive individual fitness. For each genotype pairing, the fraction of nonviable offspring was determined. Reproductive compensation then proportionately increased the number of all other genotypes from that pairing, by a total amount equal to the number of nonviable offspring multiplied by the reproductive compensation factor. For example, if nonviable genotypes result in only 50% offspring survival, then a reproductive compensation factor of 0.5 means that the cross will actually produce 75% as many offspring as a wild-type pairing, with the increased number of offspring evenly distributed amount surviving genotypes.

### Data generation

Simulations were run on the high-performance computing cluster of Peking University. All simulations were replicated a total of ten times unless otherwise specified for each parameter setting, and the results were averaged to produce points in the heatmaps. Data processing, analysis, and some figure preparation were performed in Python 3.13.2. SLiM programs, Excel files, and parameter files are available on GitHub (https://github.com/jchamper/TARE-Confinement).

## Results

### Ability of the drive systems to persist and spread in a symmetrical scenario

First, we examined our drives in simple spatial scenarios that have been studied previously for similar underdominance systems. The most straightforward is an arena in which the left half is initially occupied by drive individuals, while the right half is occupied by wild-type. In this situation, drives with thresholds under 50% tend to form waves of advance and are able to invade. Consistent with expectations, the TARE and 2-locus TARE systems were capable of forming a wave of advance unless they had significant fitness cost (Figure S1). The TARE drive lost its ability to invade when homozygote fitness was below 0.65, while the 2-locus version lost its ability to invade when its fitness (assuming that only one locus carried a fitness cost) was below 0.75, which was around 0.1 lower than the classic 2-locus underdominance system. The 1-locus 2-drive system could never form a wave of advance and retreated even without fitness costs. In all cases, higher dispersal increased the rate at which drive waves advanced or retreated.

### Effect of a population density gradient on drive spread

We next examined how increasing population density in the direction of spread (Figure 2A) could slow or halt drive waves. Such habitat heterogeneity could potentially be seen in a variety of situations as habitat transitions from one state to another. In the absence of a density gradient, all three systems produced moving fronts, but their speeds differed. TARE advanced fastest, followed by 2-locus TARE. For 1-locus 2-drive TARE, the drive is unable to advance and instead retreats, with the wild-type wave replacing the drive (Figure 2B). Increasingly steep density gradients reduced wave speed for all these systems. The estimated gradient magnitude at which propagation halted was approximately 5.2 for TARE, 4.4 for 1-locus 2-drive TARE, and 4.0 for 2-locus TARE.

**Figure 2.**
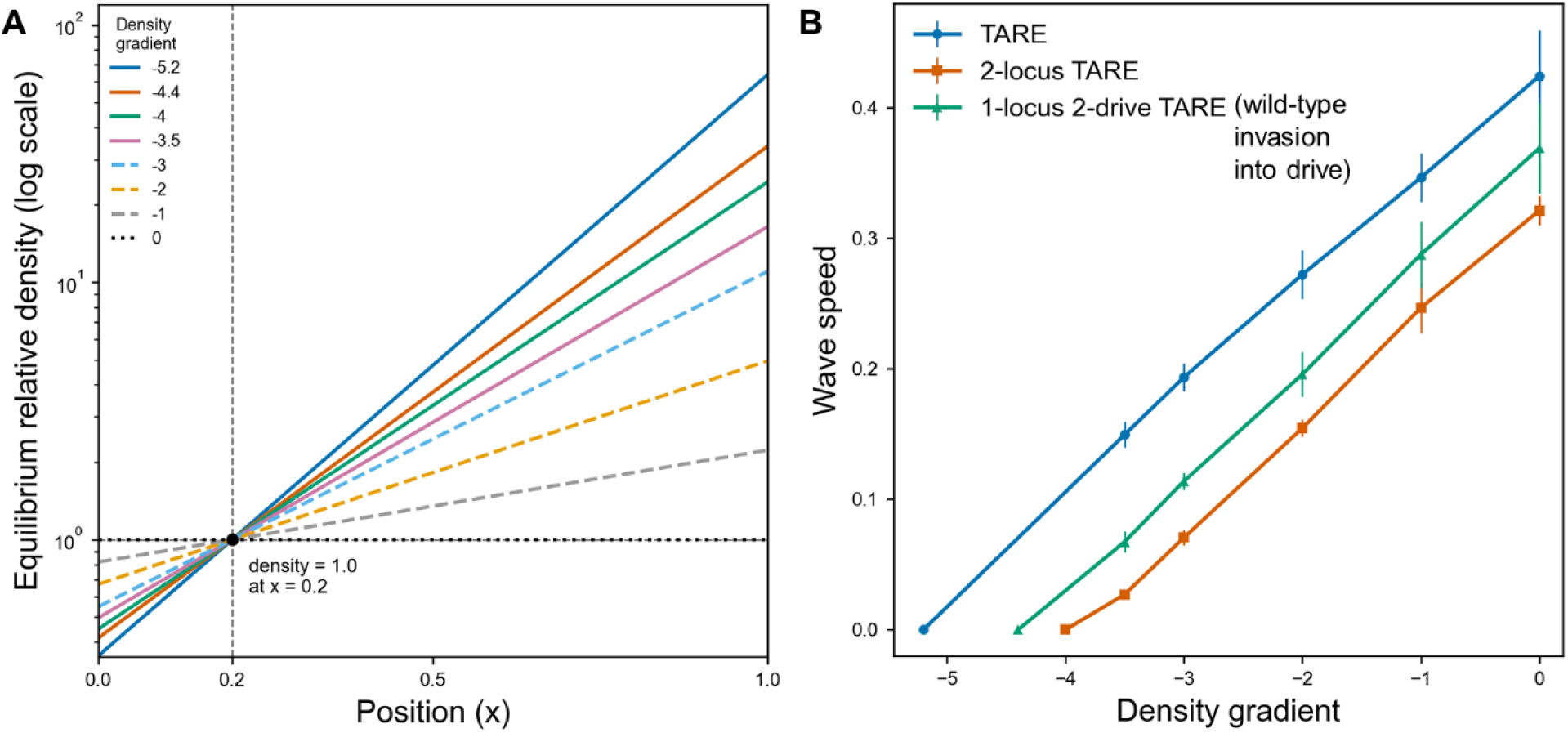
Density gradient scenario. (**A**) Chart showing equilibrium relative density as a function of position. Each line shows one tested density gradient value, with relative density normalized to 1.0 (default carrying density) at position x = 0.2. (**B**) Mean wave speed relative to average dispersal distance across density-gradient values for TARE, 2-locus TARE, and 1-locus 2-drive TARE. For 1-locus 2-drive TARE, the wave speed shows the rate that wild-type moves into drive, rather than vice versa. Points show means across 10 replicate simulations, and error bars show standard deviations.

We then tested a sharper density transition in which half of the habitat had a higher carrying density than the other half (Figure 3A). This is more representative of a sudden habitat change, such as at a beach or urban area. Increased dispersal is expected to facilitate mixing here across the step height and thus affect the outcome. We thus varied it together with the density step height. In general, higher density steps and lower migration reduced the ability of the drive to cross the step (or wild-type to invade the 1-locus 2-drive TARE), completely preventing it in some cases (Figure 3B-D). TARE was the most tolerant of the step, with high success at even a step height of 7, with success in some replicates at even higher step heights (Figure 3B). In contrast, 2-locus TARE was restricted to a much narrower range, with success declining steeply between density step heights of 2.6 and 3.0 (Figure 3C). For 1-locus 2-drive TARE in the opposite orientation, the relevant outcome was drive loss at low step heights, but larger step heights above 2.8 usually allowed the drive to persist on the high-density side (Figure 3D).

**Figure 3.**
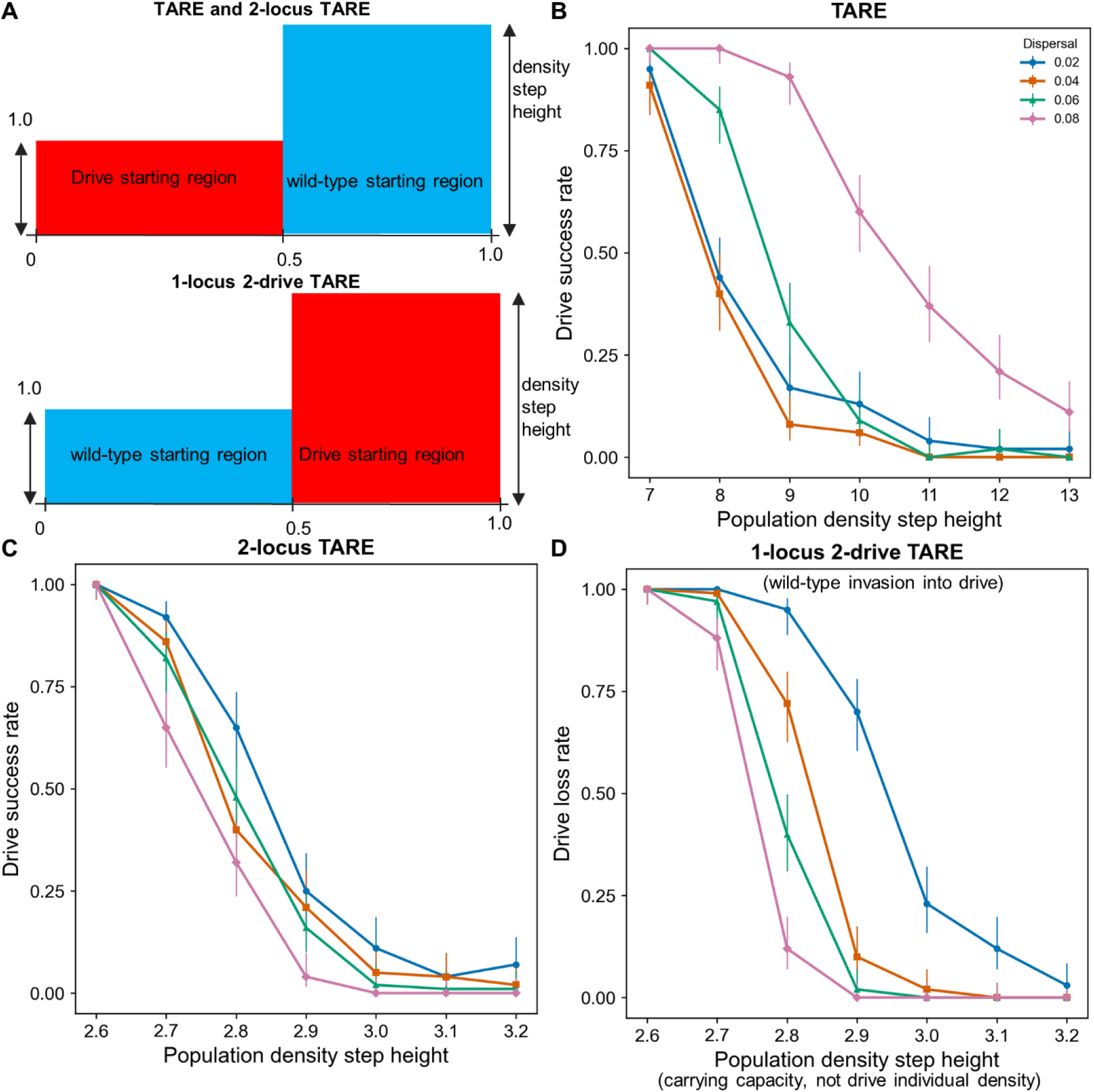
Population density step scenario. (**A**) Schematic of density step simulations. A density of “1” is the default carrying density. (**B**) Drive success rate for TARE across density step heights and dispersal values. (**C**) Drive success rate for 2-locus TARE. (**D**) Drive loss rate for 1-locus 2-drive TARE. Points show 100 replicates. Error bars show 95% Wilson confidence intervals calculated by treating each replicate as a binary drive success/non-success outcome.

Dispersal altered the effect of the density step by increasing the level of mixing between the high- and low-density regions (drive and wild-type). Sufficiently high dispersal would thus make the density transition less abrupt and therefore easier to cross for TARE drive, especially at the highest value tested (Figure 3B). This is consistent with dispersal supplying more drive carriers across the low-to-high density boundary, partially offsetting the demographic disadvantage of moving into a denser region. Because TARE drive alleles are not removed, such increased mixing is always beneficial for the drive. For 2-locus TARE, increasing dispersal did not have the same effect and in fact slightly reduced success rates. In general, higher dispersal increases drive wave speed. This result therefore suggests that the higher threshold of this mechanism made it more vulnerable to dilution when drive individuals crossed the density boundary, resulting in increased drive removal. Usually increased mixing favors the drive, but here, the density gradient made this less favorable because more wild-type alleles were brought in from the high-density area to the point of contact. For 1-locus 2-drive TARE, where the drive began on the high-density side, higher dispersal reduced drive loss for similar reasons. These patterns suggest that the effect of dispersal depends on the interaction between migrant number, local density after crossing the step, and the effective invasion threshold of each drive mechanism.

### Ability of the drive systems to persist and spread via a migration corridor

We considered a scenario in which two spatially continuous demes are connected by a migration corridor, which previously allowed some drives to be confined to one of the populations^33,35^. The two demes are assumed to be circular, each with a radius of 0.4. They are connected by a migration corridor with a variable width. All individuals in the left half of the total arena (including the left side of the migration corridor) started as drive individuals, while those on the right were wild-type. Under these conditions, a normal non-driving allele diffused, but only moderately, even for high dispersal and corridor width (Figure S2). The TARE drive and, to a lesser extent, 2-locus TARE drive were able to invade if the corridor was sufficiently wide (Figure 4). High dispersal increased the rate of invasion, but if it was above a critical level (which was proportional to corridor width), they would lose their ability to invade and remain confined to the left population. However, sufficiently high dispersal for a given corridor width could reduce their invasiveness and keep the drive confined. These same factors of low corridor width and high dispersal were essential for the 1-locus 2-drive TARE system to be able to persist in the left population.

**Figure 4.**
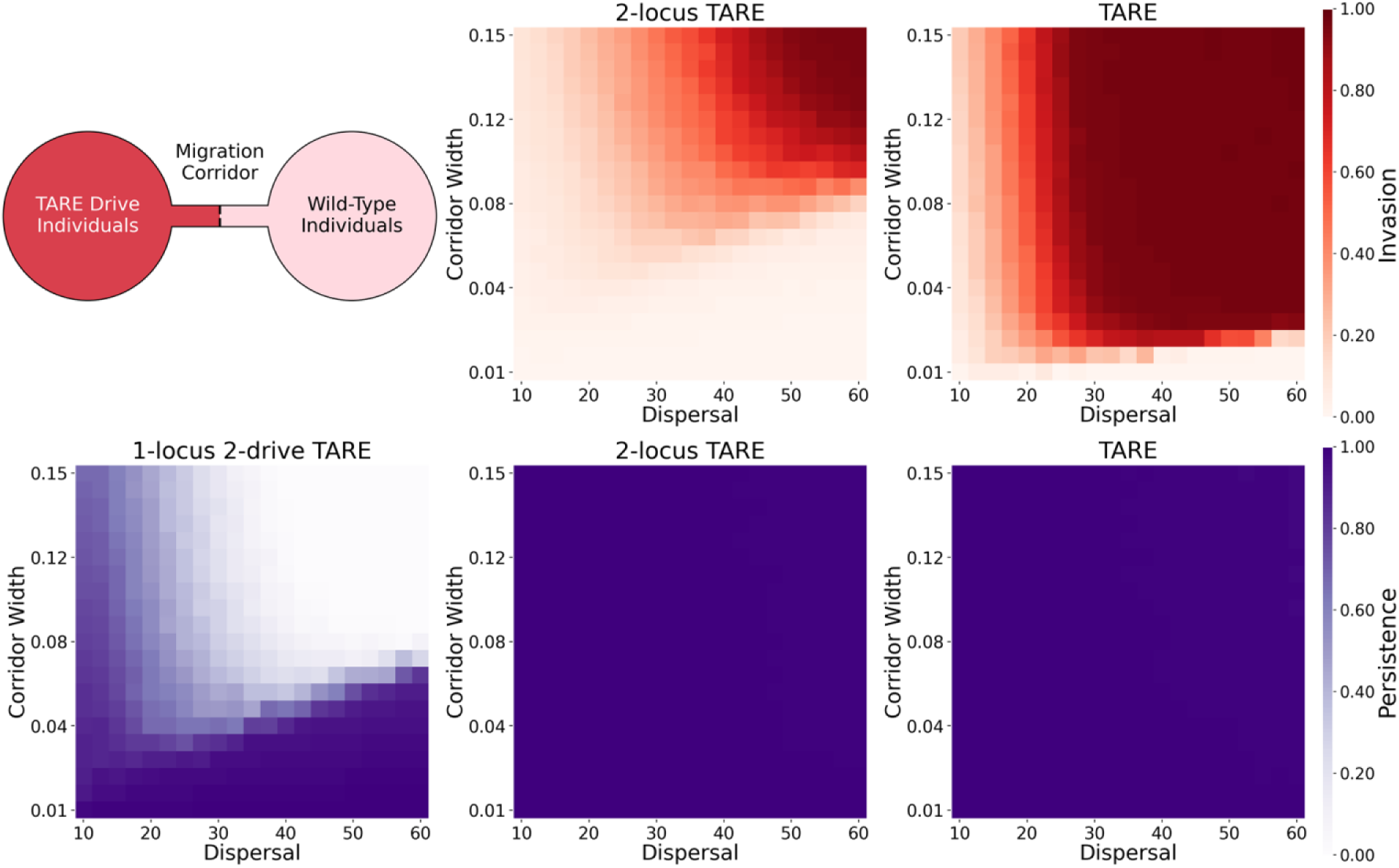
Migration corridor scenario. In an arena with two circular populations connected by a corridor, the left side starts with drive individuals (with a homozygote fitness of 0.95), and the right side starts with wild-type alleles. The simulation was run for 100 generations. Heatmaps show the final frequency of the drive in the right half (invasion) and the left half (persistence) of the arena. The 1-locus 2-drive TARE system could not invade under any parameter combination. Each point is the average of 10 simulations.

### Drive establishment after a single circular release

Aside from their ability to spread and persist in a natural environment, an important question for frequency-dependent systems is whether they can establish after a drive release in the first place. To assess this, we released all of the drives in a circle at the center of the arena (Figure S3). Compared to a symmetric situation (with drive on the left and wild-type on the right), this puts the drive at a disadvantage. At points along the edge of the circle, there will be more wild-type individuals migrating inward than drive individuals migrating outward, reducing the local drive frequency and thus requiring more drive strength for the drive to be able to succeed. We found that even the TARE drive could not substantially expand with a fitness lower than 0.8, and only persist under these conditions when dispersal was very low. The 2-locus TARE drive could only expand with nearly no fitness costs, though it could persist at lower fitness with low dispersal. The 1-locus 2-drive system was unable to persist under any parameter setting. For a weak system like this, persistence in a central release area is only possible if it has higher density than the surrounding areas.

### Ability of the drive systems to persist and spread from an edge release (port)

In panmictic models, confinement is often studied by allowing a fixed migration rate between two separate populations. This can be realistic if migrants are distributed evenly in the new population. However, when considering invasive species in particular, migrants may arrive at a point source within a spatially structured population. For example, this could be a port, with invasives arriving by ship. This could allow invaders to reach greater local frequencies, potentially facilitating invasion. To study this, we used a square area with drive individuals arriving at one of the edges of the arena, distributed within their average dispersal distance. Because the source population is unlikely to disappear, we assumed that drive individuals would continue to arrive, though the interval would vary. We also varied the number of drive individuals that arrive in each batch up to a maximum of 500, which was four times the local population size (thus putting all individuals in the release area at a disadvantage due to overcrowding - for this reason, release of additional drive individuals beyond this number would not be expected to substantially alter outcomes). Because the release radius is significantly smaller than the rest of the arena in this scenario, we only recorded the final drive rate for the whole arena to assess the ability of the drive to spread.

For the normal allele, invasion ability was negligible, even at high release sizes with frequent releases (Figure 5). The small fitness cost prevented effective establishment. Similarly, the high threshold 1-locus 2-drive TARE system could not establish even with high levels of repeated releases. For the 2-locus TARE drive, successful establishment and spread in the population was possible. However, it required a release size of at least 100 per generation, even if releases occurred every generation. If the interval between releases was three generations, establishment was not possible due to the drive’s moderate introduction threshold. Even if the local frequency of the drive was temporarily well above the threshold, the small size of the release area would mean that influx of wild-type from outside would quickly reduce the drive below its introduction threshold. For the standard TARE drive, though, establishment and invasion were possible. An almost linear relationship was found for the boundary between success and failure based on release size and interval. A release size as low as 5 could result in successful invasion if releases took place every generation, though with a release interval of 20, even a release size of 50 was insufficient to allow reliable invasion within several hundred generations. With a single release, even a size of 100 did not allow a successful invasion. This shows that the standard TARE drive is substantially more invasive than other forms, but not nearly as much as a homing drive, which could easily establish and spread with a single, small release as long as it was not subject to stochastic loss, even if configured for population suppression^54^.

**Figure 5.**
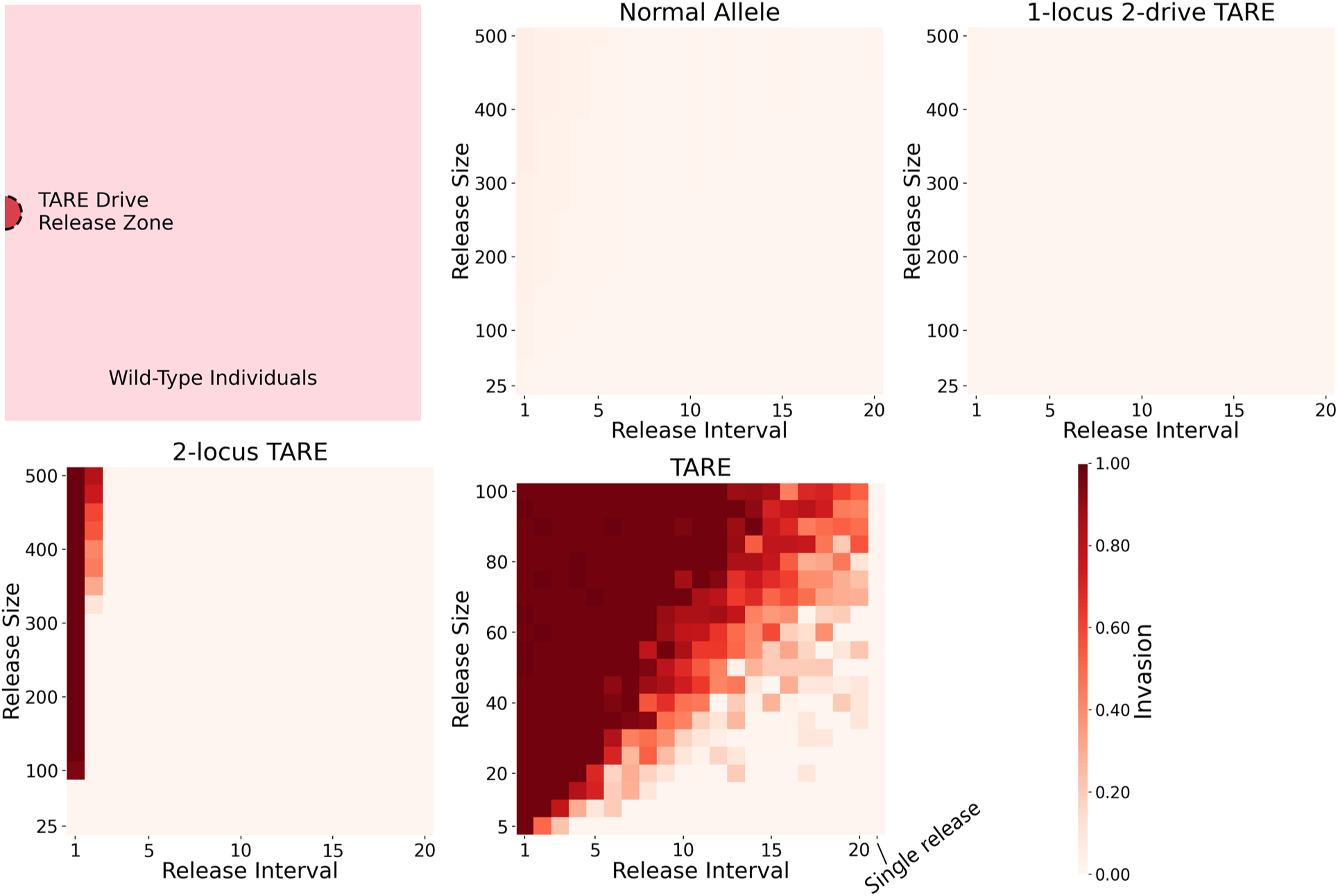
Point release scenario. In a square arena, drive individuals (with a homozygote fitness of 0.95) are released in a point in the left side of the arena with a radius equal to the average dispersal distance. The simulation was run for 300 generations. Heatmaps show the final frequency of the drive. Release Interval refers to the time between releases (a release interval of 1 means that new drive individuals are released in every generation), and Release Size is the number of individuals in each release. In the last column for TARE drive, we assume a single release (with no subsequent releases). Each point is the average of 10 simulations.

### Effect of reproductive compensation on drive thresholds

Finally, we modeled a scenario involving parental care within a limited offspring population. Because parental care enhances the fitness of offspring carrying the drive, the drive requires a lower introduction threshold and becomes more invasive. This is an important consideration, as this factor could affect the confinement of gene drive systems. To assess this, we measured how the compensation factor associated with parental care affects the TARE drive performance.

We found that as the compensation factor increases, the introduction threshold across all fitness levels decreases (Figure 6A), with the drive able to spread far more rapidly at low frequency compared to standard TARE drive without reproductive compensation. At high frequency, though, the drive has similar dynamics regardless of reproductive compensation because most offspring are viable. Thus, an increase in the reproductive compensation factor does not substantially affect the final drive equilibrium frequency (Figure 6B). Indeed, the equilibrium frequency actually slightly decreases because the number of drive/disrupted allele heterozygotes is increased in rare pairings that produce nonviable offspring. Overall, these findings demonstrate that reproductive compensation derived from parental care or other mechanisms lowers the introduction threshold frequency, making TARE drive significantly more invasive.

**Figure 6.**
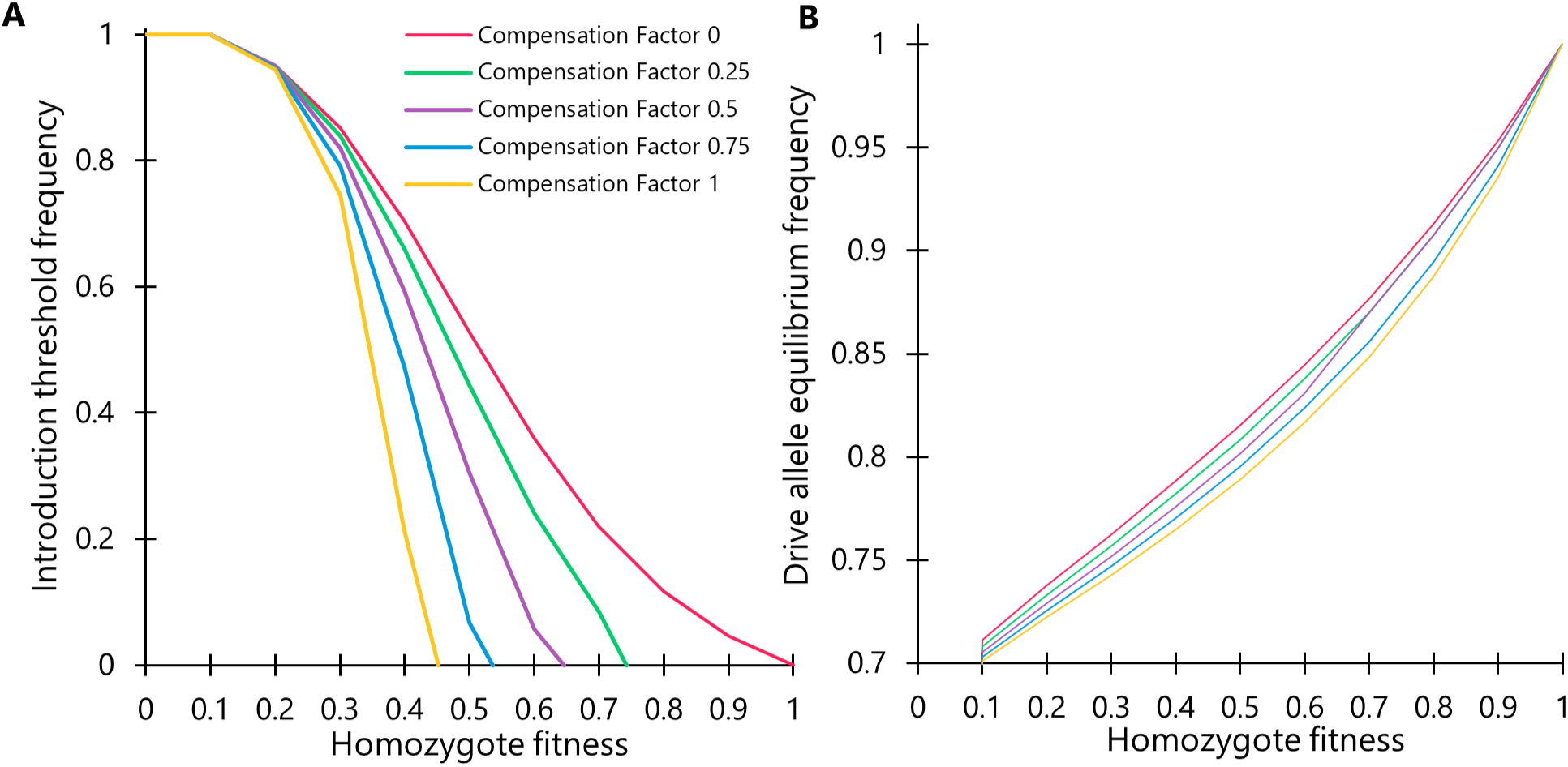
Properties of TARE drive with reproductive compensation. Using a deterministic, discrete-generation model, reproductive compensation was implemented, representing an increase in the number of generated offspring for each offspring with a nonviable genotype from the same parent. **(A)** Introduction threshold frequency (with release of homozygous individuals) with varying drive homozygote fitness value and reproductive compensation factor. **(B)** The TARE drive allele equilibrium frequency as a function of homozygote fitness value and reproductive compensation factor. Note that final alleles are composed entirely of drive and disrupted alleles, so the final drive carrier frequency is always 100%.

## Discussion

In this study, we investigated three of the latest CRISPR toxin-antidote designs. One promising aspect of these drives is their potential to be confined to a target population. Their frequency dependence also means that deployment of such drives must be carefully considered to ensure success. We have shown in this study that spatial factors can influence these considerations, and that confinement and persistence in connected populations is not always guaranteed. Further, we have shown that the system dynamics of parental care and resource investment can reduce the confinement level of all of these systems, emphasizing the need to consider a diverse set of ecological factors when assessing confined gene drive deployment.

Several aspects of this study are similar to our previous study that examined classic underdominance drive systems based on RNAi or other mechanisms^33^. These systems had qualitatively similar behavior, particularly the 2-drive systems that could occupy one or two separate loci. However, these have proven difficult to further develop in organisms of interest. In this regard, CRISPR toxin-antidote systems are likely superior, capable of using off-the-shelf components, and having been quickly demonstrated in not just *Drosophila*^19–21,23^, but also plants^27,28^ and mosquitoes^26^. We thus updated several scenarios from our previous study with these new systems, which have somewhat lower thresholds than their older non-CRISPR counterparts. We also added the standard TARE drive, which has a substantially lower threshold than these older systems, analogous to *Medea*^55^, which was not considered in our previous spatial model. Developing quantitatively accurate models for this will allow for more precise predictions of actual drives that may be constructed in the near future.

We considered new scenarios, in particular density gradients and a single-origin release. The first is expected to be a common occurrence in natural environments with heterogeneous habitat. Moving up a density gradient could substantially slow a drive, though we found that the gradient needs to be fairly steep for a drive to stop advancing. Similarly, the 1-locus 2-drive version can only persist with a steep gradient. Habitat gradients in real-world situations may also take the form of sudden, rather than gradual transitions. In this case, we found that the high dispersal ability of individuals could also slightly increase the success rate of invasions for TARE, but it may actually serve to slightly reduce the ability of other invasive systems with higher introduction thresholds to cross the density step.

A likely explanation is that dispersal has two opposing effects when a drive encounters a density step. On one hand, higher dispersal can increase the absolute number of drive carriers that cross the low-to-high density boundary, promoting spread into the denser region. On the other hand, higher dispersal can also spread those migrants over a broader area, reducing their local concentration and making it more difficult for the drive frequency in any small area to exceed the effective invasion threshold. The outcome should therefore depend on both dispersal and step height. A higher target-side density increases the number of resident wild-type individuals that dilute incoming drive carriers, whereas a lower density or a larger migrant pulse can allow the local post-dispersal drive frequency to exceed the threshold. This may explain why dispersal facilitated spread for TARE, but reduced success for 2-locus TARE (and for wild-type invading 1-locus 2-drive TARE), which has a higher introduction threshold.

The single-origin release may be particularly relevant to many situations in which a gene drive is deployed. It would usually represent a shipping port, or perhaps even an airport for some types of insects. In this case, the drive would be at a substantial advantage compared to a panmictic model, with substantially higher local frequency. This allowed the TARE drive to become invasive with relatively modest but regular release sizes (a single release of a small number of individuals would thus not be sufficient for spread of a TARE drive, unlike a homing drive). However, the 2-locus TARE drive was substantially more robust to this, requiring an influx of large numbers of new individuals every 1-2 generations, which should be substantially less likely in many scenarios.

CRISPR toxin-antidote gene drives, together with many other frequency-dependent drive types, often result in offspring nonviability at early developmental stages. Most modeling, even spatial modeling (where offspring tend to immediately spread out), tends to assume no additional effects. However, with fewer offspring, parental care, which can include *in utero* investment, will tend to provide a greater amount of resources for each remaining offspring. This is likely to increase their fitness, and because these remaining offspring will tend to carry the drive, it will strengthen the drive and reduce its introduction threshold. The exact level at which this occurs could vary greatly with species ecology and life history. In mammals, such as invasive predators in New Zealand, this may have a noticeable effect. In insects, the effect is likely smaller, though it could still be present if, for example, a female lays most of her eggs in one location in which other eggs are not present at high density. Offspring nonviability may therefore reduce sibling competitive pressure on drive individuals and significantly increase survival. We showed that theoretically, this effect could substantially alter drive invasiveness characteristics. This is not necessarily unfavorable for efficient drive spread, but it does require consideration when assessing whether a drive has the requisite level of confinement.

Overall, our study indicates that CRISPR toxin-antidote drives are promising candidates for confined modification of target populations. Standard TARE drive can spread widely, but still remain regionally confined. 2-locus TARE will yield strong local spread, but obstacles even within connected populations could still confine it. 1-locus 2-drive TARE cannot invade under normal circumstances and would require widespread releases, and even then, would tend to be eliminated if connected to a wild-type population without a major barrier. Overall, these spatial and life history considerations will need to be taken into account when determining optimal deployment of gene drives and whether a particular gene drive is most suitable for a certain task.

## Acknowledgements

This study was supported by the Center for Life Sciences and the National Natural Science Foundation of China (32270672, W2432018).

## Supplemental Information

**Figure S1.**
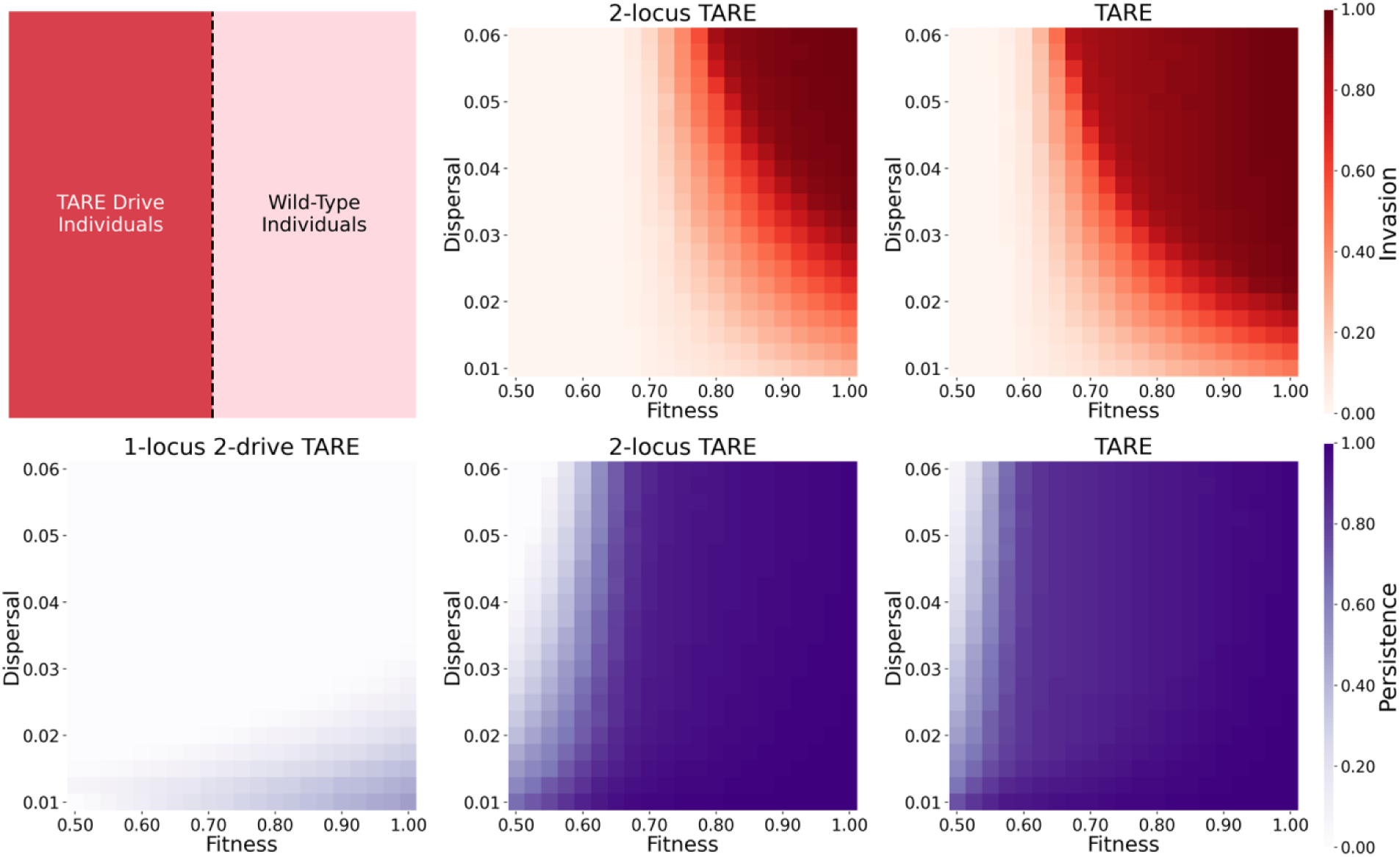
Symmetric drive wave scenario. In a square arena, the left side starts with drive individuals, and the right side starts with wild-type alleles. The simulation was run for 60 generations. Heatmaps show the final frequency of the drive in the right half (invasion) and the left half (persistence) of the arena. The 1-locus 2-drive TARE system could not invade under any parameter combination. Each point is the average of 10 simulations.

**Figure S2.**
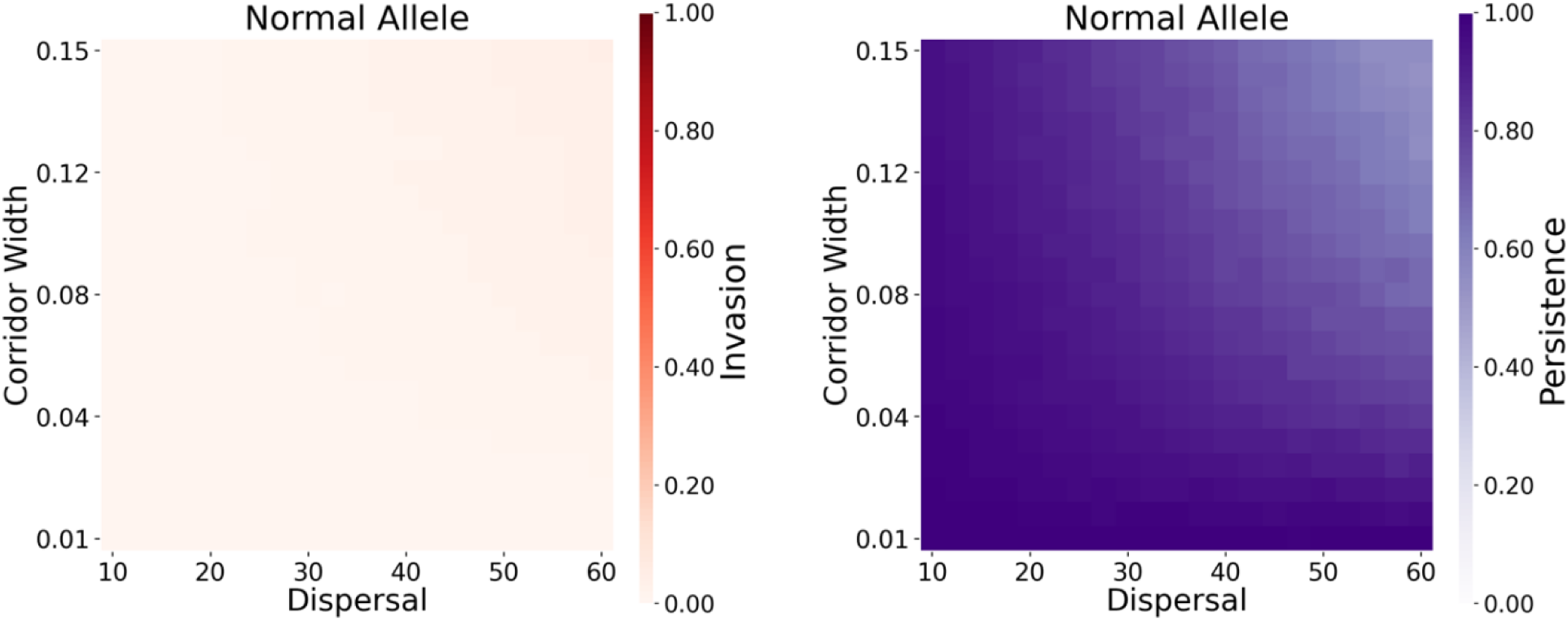
Non-drive allele in migration corridor. In an arena with two circular populations connected by a corridor, the left side starts with individuals homozygous for a non-drive allele (with a homozygote fitness of 0.95), and the right side starts with wild-type alleles (see Figure 4). The simulation was run for 100 generations. Heatmaps show the final frequency of the non-drive allele in the right half (invasion) and the left half (persistence) of the arena. Each point is the average of 10 simulations.

**Figure S3.**
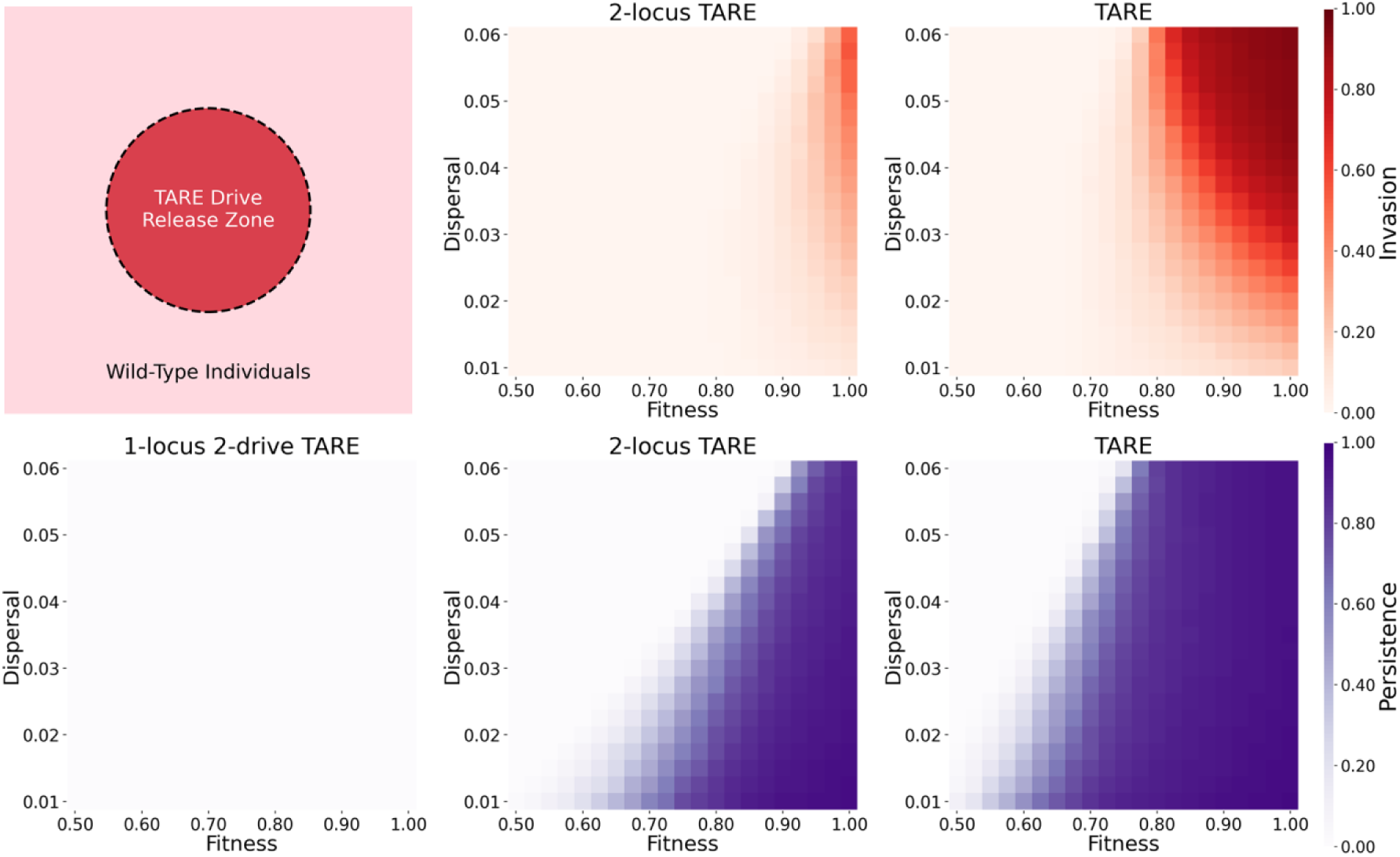
Circular release scenario. In a square arena, drive individuals were released in a circle of radius 0.25 at 80% initial frequency in the release area. The simulation was run for 30 generations. Heatmaps show the final frequency of the drive outside the release circle (invasion) and within the release circle (persistence). The 1-locus 2-drive TARE system could not invade or persist under any parameter combination. Each point is the average of 10 simulations.

